# *Pus7* mutation links tRNA dysregulation to aggressive behavior through activation of the integrated stress response and glycolytic reprogramming

**DOI:** 10.1101/2025.05.12.653498

**Authors:** Michael Stock, Maria Guillen Angel, Monika Witzenberger, Jakub Nowak, Anna Biela, Andrzej Chramiec-Glabik, Vittoria Mariano, René Dreos, Athéna Sklias, Manfredo Quadroni, Anneke Brümmer, Hector Gallart-Ayala, Julijana Ivanisevic, Nicolas Guex, Leon Kleenman, Sebastien Leidel, Mark Helm, Claudia Bagni, Sebastian Glatt, David Gatfield, Schraga Schwartz, Jean-Yves Roignant

**Affiliations:** Center for Integrative Genomics, Faculty of Biology and Medicine, University of Lausanne, 1015 Lausanne, Switzerland; Department of Molecular Genetics, Weizmann Institute of Science, 7610001 Rehovot, Israel; Malopolska Centre of Biotechnology, Jagiellonian University, Krakow, Poland; Department of Fundamental Neurosciences (DNF), University of Lausanne, 1011 Lausanne, Switzerland; Proteomics Core Facility, University of Lausanne, 1015 Lausanne, Switzerland; Bioinformatics Competence Center, University of Lausanne, 1015 Lausanne, Switzerland; Metabolomics Platform, Faculty of Biology and Medicine, University of Lausanne, 1005 Lausanne, Switzerland; Department of Chemistry, Biochemistry and Pharmaceutical Sciences, University of Bern, 3012 Bern, Switzerland; Institute of Pharmaceutical and Biomedical Sciences, Johannes Gutenberg-University Mainz, 55128 Mainz, Germany; Department of Biomedicine and Prevention, University of Rome Tor Vergata, 00133 Rome, Italy; University of Veterinary Medicine, 1210 Vienna, Austria

**Keywords:** Pseudouridine, Pus7, aggression, RNA modification, translation

## Abstract

Pseudouridine (Ψ) is a prevalent RNA modification found in multiple RNA species. It is deposited by Ψ synthases (Pus) and it can stabilize RNA structures. Patients carrying alterations in the *Pus7* gene suffer from developmental delay, intellectual disability, microcephaly, hyperactivity and increased aggression levels. Here we show that the *Pus7* mutation in human patient cells and in a *Drosophila* model is associated with a specific decrease of tRNA:Aspartate (tRNA-Asp) levels, which leads to slow decoding at Aspartate codons. This in turn activates the integrated stress response and induces a metabolic shift towards increased glycolysis and reduced mitochondrial respiration. Elevating tRNA-Asp expression, inhibiting the integrated stress response or dampening the glycolytic pathway is sufficient to rescue the aggressiveness phenotype, demonstrating the involvement of the tRNA-Asp-ISR-glycolysis axis in this behavior. Together our data provide new insights into the molecular defects associated with the loss of Pus7 and suggest potential new avenues for therapeutic treatment.

## Introduction

Chemical modifications of RNA molecules are prevalent in all kingdoms of life and have the potential to alter gene expression. Aberrant deposition of RNA modifications has been linked to various physiological and pathological defects. Notably, an increasing number of predicted human transfer RNA (tRNA) modification genes have been associated with neurological disorders, including microcephaly and intellectual disability [1], [2]. Yet, it is still unclear how aberrant modifications on tRNA leads to the pathogenesis, and if other RNA substrates may be implicated.

Ψ is the most abundant RNA modification, and is present across all three domains of life. It is found in most RNA species [3], including ribosomal RNA (rRNA), transfer RNA (tRNA), small nuclear RNA (snRNA), small nucleolar RNA (snoRNA) and messenger RNA (mRNA). RNA pseudouridylation stabilizes RNA by providing an extra hydrogen bond, increasing the rigidity of the backbone [4], [5], [6]. The formation of Ψ is catalyzed by Ψ synthases (PUS) either in a guide RNA dependent manner by the box H/ACA ribonucleoprotein complex or by standalone PUS enzymes [7], [8]. Notably, all PUS enzymes share a conserved aspartate which is essential for their catalytic activity [9].

Defects in pseudouridylation have been associated with several diseases [10]. For instance, patients with recessive defective variants of *PUS1* feature mitochondrial myopathy, lactic acidosis and sideroblastic anemia (MLASA) as well as failure to thrive, developmental delay and intellectual disability [11], [12]. Furthermore, patients with defects in *PUS3* feature a more brain-centric phenotype consisting of intellectual disability, microcephaly, growth delay, short stature, failure to thrive, and seizures [13], [14]. Likewise, we and others have reported on the deleterious effects of *PUS7* mutations in human consisting of intellectual disability, speech delay, microcephaly, short stature, and aggressive behavior [15], [16], [17], [18], [19], [20]. Aberrant levels of PUS7 have also been linked to myelodysplastic syndrome, glioblastoma, colorectal and lung cancer [21], [22], [23], [24], [25], [26].

Previously, we established a *Drosophila* model in which Pus7 function was impaired. We demonstrated that its loss in this organism leads to hyperactivity, aggression, as well as an orientation defect [16], associated with reduced pseudouridylation of several tRNA species. Here, we demonstrate that the absence of Pus7 leads to a specific reduction of tRNA:Aspartate (tRNA-Asp) levels, resulting in slower translation at Aspartate codons and ribosomal disome formation. These events activate the integrated stress response (ISR), which results in increased glycolysis and reduced mitochondrial respiration.

Notably, overexpressing tRNA-Asp rescues both, mitochondrial respiration as well as aggression in *Pus7*^*fs*^ flies, indicating that tRNA-Asp is a key target in the mechanism of disease. We show that this target is also strongly and specifically altered in human patient cells, suggesting a conserved mechanism. Lastly, we demonstrate that preventing the activation of the ISR, or reducing glycolytic activity in *Pus7* mutant flies is sufficient to restore normal aggression level. Altogether these data indicate that lack of Pus7 destabilizes a specific tRNA leading to changes in carbohydrate metabolism, ultimately impacting behavior.

## Results

### *Pus7* mutation leads to aggressive behavior and reduced tRNA-Asp levels

We previously reported increased aggressive behavior in *PUS7* mutant patients as well as in a fly model for Pus7 loss [16]. We repeated the aggression assay in the mutant flies to establish a system for phenotypic rescue, focusing on strong aggressive behaviors such as lunging and tussling. The increased aggression level associated with the mutant could be rescued by overexpression of Pus7 cDNA in the whole fly (**Fig. 1A**). Likewise, expressing Pus7 only in neurons via the *elav* promoter was sufficient to restore the wild type phenotype, demonstrating the neuronal role of Pus7 in this function. Furthermore, a catalytically dead version of Pus7 with the conserved aspartate mutated to glutamate [6], was unable to rescue the aggressiveness, confirming the role of Ψ modification in this behavior (**Fig. 1A**).

**Figure 1.**
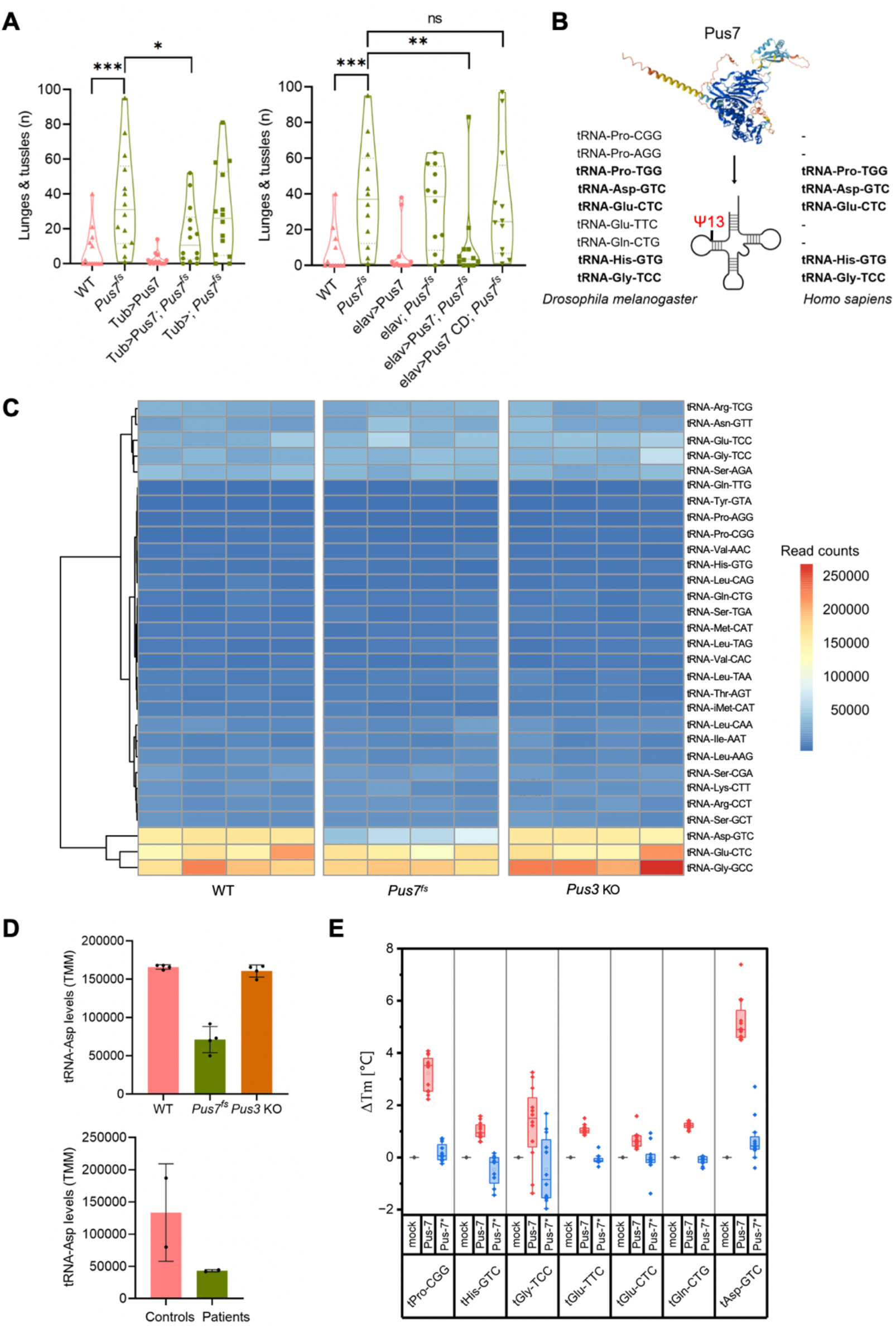
*Pus7* mutation in flies leads to aggressiveness and decrease in tRNA-Asp levels. **A**. Violin plot showing the aggression levels (number of lunges and tussles) in WT (n = 15), *Pus7*^*fs*^ (n = 15), tub>Pus7 (n = 15), tub>Pus7; *Pus7*^*fs*^ (n = 15), and tub>; *Pus7*^*fs*^ (n = 15) flies (Mann-Whitney test, P value < 0.001 ***, P value < 0.05 *) (left) and Violin plot showing the aggression levels (number of lunges and tussles) in WT (n = 15), *Pus7*^*fs*^ (n = 15), elav>Pus7 (n = 15), elav>; *Pus7*^*fs*^ (n = 15), elav>Pus7; *Pus7*^*fs*^ (n = 15) and elav>Pus7 CD; *Pus7*^*fs*^ (n = 15) flies (Mann-Whitney test, P value < 0.001 ***, P value < 0.01 **, P value > 0.05 ns) (right). **B**. Schematic showing the tRNA targets of Pus7 in humans and flies found by Psi-seq. **C**. Heatmap showing normalized read counts (TMM) for tRNAs measured using the input samples (non CMC treated) from Psi-Seq. WT (n = 4), *Pus7*^*fs*^ (n = 4) and *Pus3* KO (n = 4) flies. **D**. Bar plot showing normalized counts (TTM) for tRNA-Asp in WT (n = 4), *Pus7*^*fs*^ (n = 4) and *Pus3* KO (n = 4) flies (top) measured by Psi-Seq and bar plot showing normalized counts (TTM) for tRNA-Asp in patients (n = 4) and controls (n = 2) measured by Psi-Seq (bottom). **E**. Box plots showing relative melting temperature of tRNAs incubated with mock, PUS7 (red) and mutated PUS7 (Pus7*, blue). Boxes represent 25-75% of the data and whiskers extend to 1.5 times the interquartile range. Medians are shown as lines, and means as squares. Individual data points (n=4) are shown as dots. n.d. (not determined).

We found that nine tRNA species (tRNA-Pro-[CGG-, AGG, TGG], tRNA-Asp-GTC, tRNA-Glu-[TCC, CTC], tRNA-Gln-[CTG, CTG], and tRNA-Gly-TCC) showed significant loss of pseudouridylation at position 13 in *Pus7*^*fs*^ flies [16]. Of these, five are also PUS7 targets in patient lymphoblastoid cell lines, indicating that Pus7 tRNA targets are conserved from flies to human (**Fig. 1B**). Intriguingly, closer examination of the sequencing data indicated a specific reduction in tRNA-Asp-GTC in *Pus7*^*fs*^ flies compared to WT flies, an effect which can also be observed in human patients (**Fig. 1C-D**).

Since the high frequency of post-transcriptional modifications on tRNAs can introduce biases in reverse transcription-based quantification methods, we sought to validate this observation with orthogonal approaches. We performed microscale thermophoresis (MST), which allows the quantification of a tRNA species of interest in a tRNA mixture by detecting it with a fluorescently labeled probe via base-pairing [27]. Using this approach, we confirmed that tRNA-Asp was indeed significantly reduced in *Pus7*^*fs*^ flies and that overexpression of *Pus7* cDNA in the whole fly rescued tRNA-Asp levels back to wild type (**Fig. S1A**). Likewise, Northern blot analysis confirmed reduced tRNA-Asp in *Pus7*^*fs*^ flies (**Fig. S1B**).

RNA modifications in the structural core of tRNAs have been shown to affect their stability and folding [6], [28]. Position 13 of tRNA is the last nucleotide of the stem of the D-loop and we hypothesized that the lack of pseudouridylation at this position is required for the stability of tRNA-Asp. To test this directly, we incubated recombinant human PUS7 with *in vitro* transcribed Drosophila tRNA species, confirmed the specific modification and quantified the changes in stability upon an increased temperature gradient *in vitro* (**Fig. 1E**). Indeed, all tested PUS7-target tRNAs showed an increased thermostability after modification by PUS7, but no stabilization after incubation with an enzymatically inactive variant of PUS7 was observed. Foremost, tRNA-Asp-GTC shows by far the largest increase in melting point temperature following incubation with PUS7, while other tRNAs were less affected. Hence, modification by PUS7 seems to be particularly relevant for the stability of tRNA-Asp-GTC, which is in perfect agreement with its disappearance *in vivo*. Of note, none of the other tested substrate tRNAs were significantly decreased *in vivo* (**Fig. 1C**), but at the moment we cannot exclude any additional functional roles of Ψ13 for the function of the other Pus7 target tRNAs. For tRNA-Asp-GTG, the reduced stability in the absence of Ψ13 was not accompanied with increased levels of tRNA-Asp fragments (**Fig. S1B**), suggesting that the tRNA transcripts are fully degraded *in vivo*, possibly via the rapid tRNA decay (RTD) pathway.

### Lack of Pus7 leads to changes in translation and metabolic alterations

We next addressed whether the reduction of tRNA-Asp might lead to translational defects by performing ribosome profiling in heads of *Pus7*^*fs*^ and WT flies. Detected transcripts were categorized depending on their differential regulation in *Pus7*^*fs*^ heads into those with significant difference in transcription (corresponding to significant changes in both, mRNA expression and ribosome occupancy, but not in their ratio), those with difference in translation (corresponding to significant change in translation efficiency, the ratio of ribosome occupancy to mRNA expression), and those with significant change in mRNA expression only (**Fig. 2A, B**). The majority of changes occurs at the translation level, consistent with the role of tRNA in translation. We next measured ribosome occupancy on individual codons in both conditions. Interestingly, while we observed only minor changes on most codons, ribosome occupancy on Aspartate codons was selectively increased in *Pus7*^*fs*^ flies, suggestive of slower translation at these sites (**Fig. 2C**). More specifically, this effect was associated with Asp codons located in the A-site of the ribosome, consistent with longer ribosomal dwell times when cognate tRNA levels become limiting, and it was largely independent of the codon located in the P site (**Fig. 2D**). Furthermore, transcriptome-wide analysis revealed increased ribosome occupancy in *Pus7*^*fs*^ flies in the region located 30 nt to the 5’ side of

**Figure 2.**
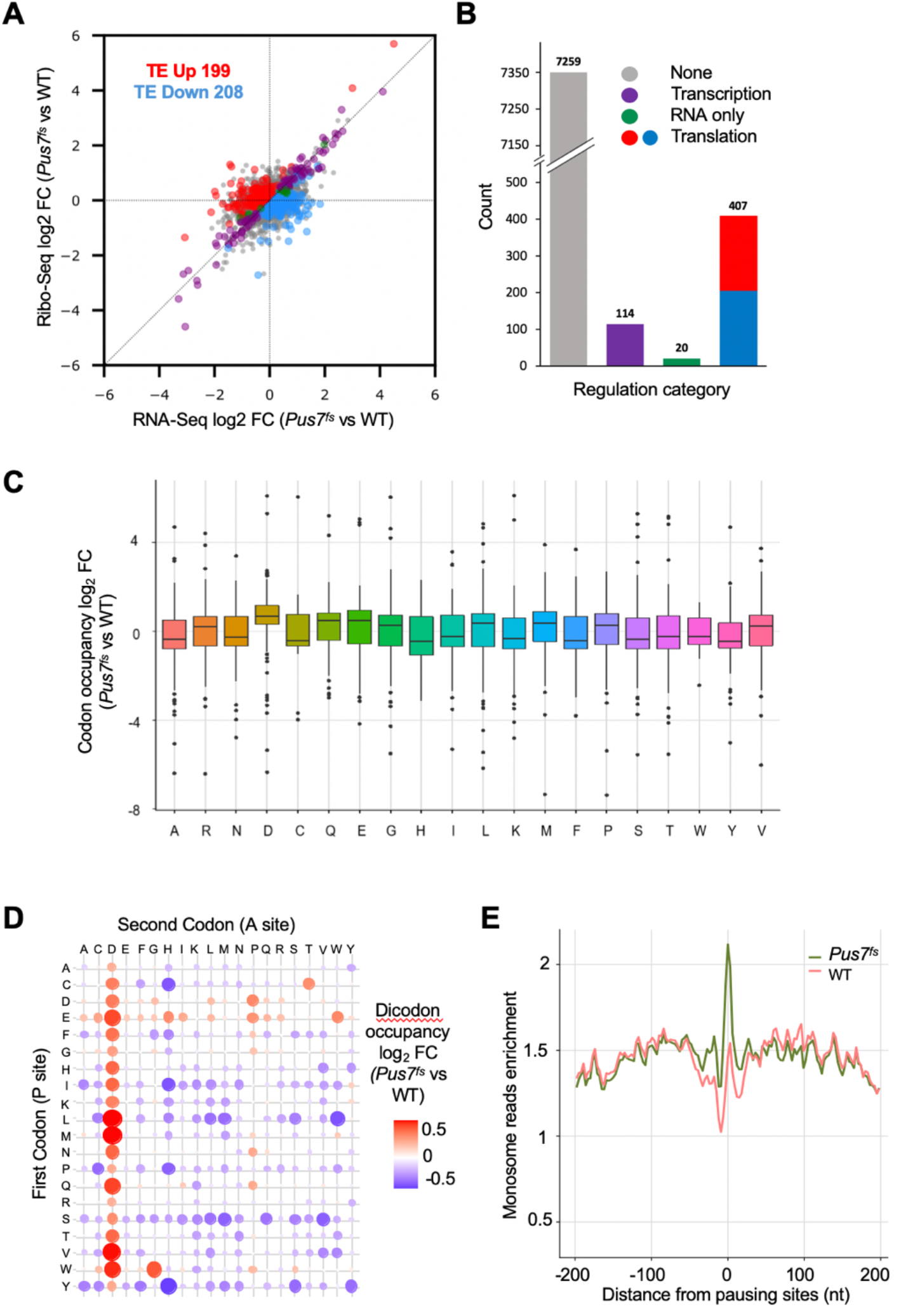
*Pus7* mutation leads to changes in translation. **A**. Scatterplot of the fold change of the WT/*Pus7* mutant at mRNA level (x-axis) and ribosome occupancy (y-axis). **B**. Bar plot giving the number of mRNA transcripts significantly regulated in each category. P value < 0.05. **C**. Box plot showing comparison of ribosome codon occupancy in the A site between *Pus7*^*fs*^ (n = 4) and WT (n = 4) fly heads. **D**. Dot plot showing ribosome occupancy at dicodons with first codon at P sites and second codon at A sites between *Pus7*^*fs*^ (n = 4) and WT (n = 4) fly heads. **E**. Monosome reads enrichment around ribosome pausing sites in *Pus7*^*fs*^ (n = 4) and WT (n = 4) fly heads.

Asp codons. This corresponds to where an upstream stacked ribosome would be positioned. Increased ribosome occupancy suggests that slow Asp decoding leads to increased ribosomal queueing (**Fig. 2E**).

To complement the ribosome profiling, we quantified protein levels on head samples from WT and *Pus7*^*fs*^ flies through mass spectrometry. Ribosome profiling and proteomics data showed overall good correlation **(Fig. S2A, B)**. Intriguingly, further analyses revealed that proteins with a role in carbohydrate metabolism were strongly enriched among the upregulated group, whereas those associated with lipid metabolism were enriched among the downregulated fraction (**Fig. 3A, B**).

**Figure 3.**
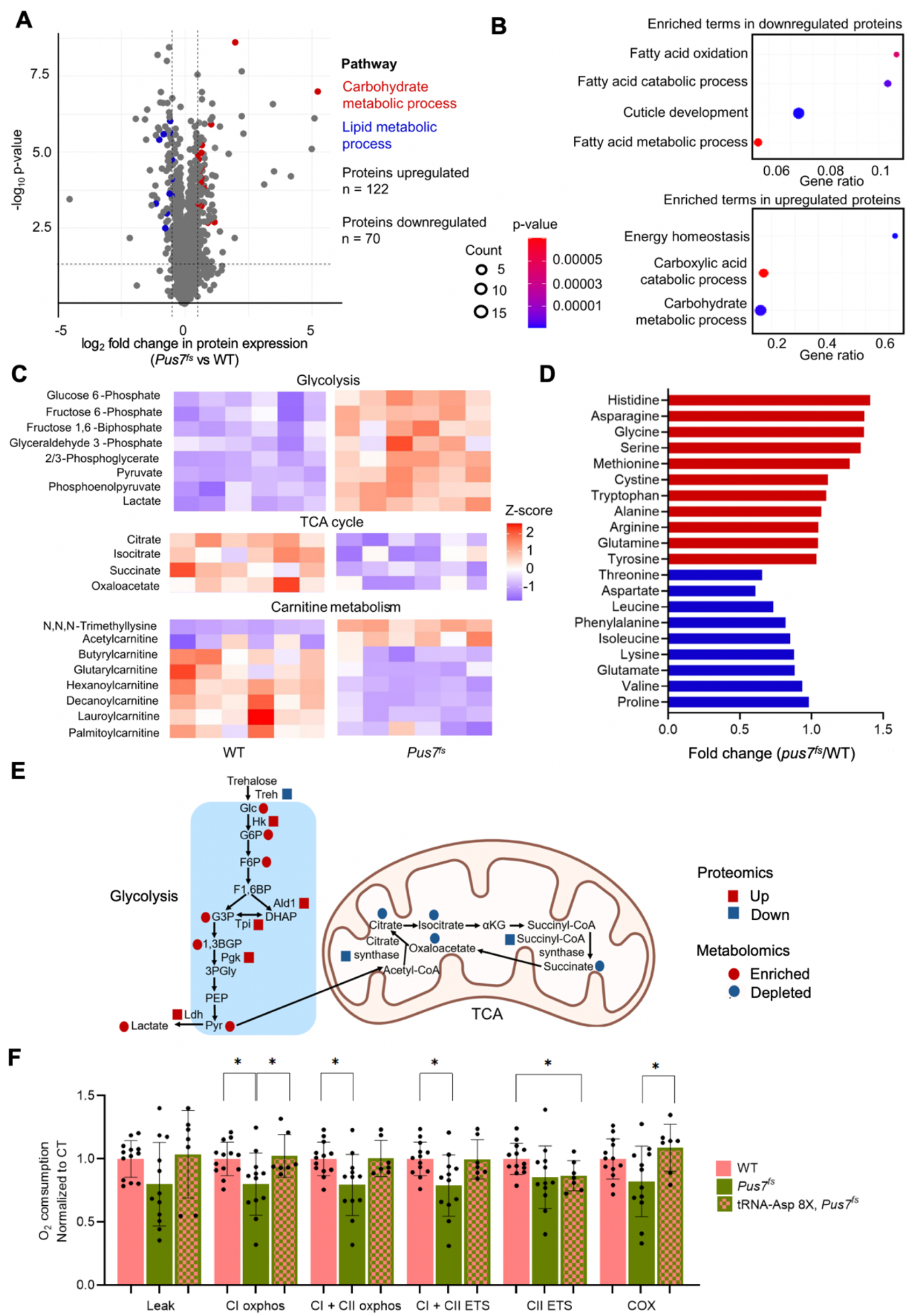
*Pus7* mutation leads to changes in metabolism. **A**. Volcano plot showing protein expression comparison between *Pus7*^*fs*^ (n = 4) and WT (n = 4) fly heads, shown in red the proteins involved in carbohydrate metabolism that are upregulated and in blue proteins involved in lipid metabolism that are downregulated. **B**. Enriched terms of differentially expressed proteins in *Pus7*^*fs*^ fly heads, downregulated (top) and upregulated (bottom) **C**. Heatmap showing comparison of metabolite levels in *Pus7*^*fs*^ vs WT. **D**. Bar plot showing comparison of amino acid levels in *Pus7*^*fs*^ vs WT. Positive fold changes are shown in red and negative fold changes in blue. **E**. Schematic of the metabolic pathways affected by *Pus7* mutation, proteins are shown in squares and metabolites in circles, upregulation in *Pus7*^*fs*^ compared to WT is shown in red and downregulation in *Pus7*^*fs*^ compared to WT is shown in blue. **F**. Bar plot showing oxygen consumption normalized to WT values in WT (n = 8) and *Pus7*^*fs*^ (n = 8) fly heads in the respiration complexes (Unpaired t-test, P value < 0.05 *) (left) and bar plot showing oxygen consumption normalized to WT values in WT (n = 8), *Pus7*^*fs*^ (n = 8) and tRNA-Asp 8X; *Pus7*^*fs*^ (n = 8) fly heads in the respiration complexes (Unpaired t-test, P value < 0.05 *) (right).

To determine whether these changes in protein levels impacted the metabolic landscape of *Pus7*^*fs*^ flies we performed metabolomics analyses using fly head samples. Consistent with the prediction from the proteomics analysis, metabolites of glycolysis were significantly enriched, whereas TCA cycle metabolites as well as various acyl-carnitines were depleted (**Fig. 3C, E**). Moreover, amino acid levels were also affected, among which aspartate derived from the TCA metabolite oxaloacetate shows a decrease in *Pus7*^*fs*^ flies (**Fig. 3D**). In contrast, levels of amino acids derived from glycolytic activity such as serine, glycine and alanine were all enriched in *Pus7*^*fs*^ flies (**Fig. 3D**).

TCA cycle downregulation at the metabolite and protein level in *Pus7*^*fs*^ flies was suggestive of impaired mitochondrial function. To test this hypothesis, we measured the oxygen consumption of heads of *Pus7*^*fs*^ and WT flies, revealing a significant decrease in complex I and II activities in mutant flies (**Fig. 3F**).

Thus, altogether these data indicate that the Pus7 loss-of-function leads to a metabolic shift towards higher glycolytic activity and lower mitochondrial respiration.

### Expressing tRNA-Asp is sufficient to rescue metabolic defects and aggressiveness

We next aimed to ameliorate the defects caused by *Pus7* mutation by overexpression of tRNA-Asp. We generated flies carrying a transgene harboring 4 copies of tRNA-Asp-GTC, present on the two homologous chromosomes in the wild type and *Pus7*^*fs*^ background, respectively. Remarkably, we found that tRNA-Asp overexpression could fully rescue the aggression phenotype in the *Pus7*^*fs*^ background (**Fig. 4A**). In contrast, overexpression of tRNA-Lys, as a control tRNA that is not a Pus7 target and whose levels are not affected upon *Pus7* mutation (**Fig. 1C**), did not restore the aggression level **(Fig. S3)**. In addition, the 8 copies of tRNA-Asp fully rescued the mitochondrial respiration defects, with the exception of CII ETS activity that remains unchanged (**Fig. 3F**). Collectively, these results indicate that tRNA-Asp is a primary target of Pus7 for the metabolic control and aggression phenotypes.

**Figure 4.**
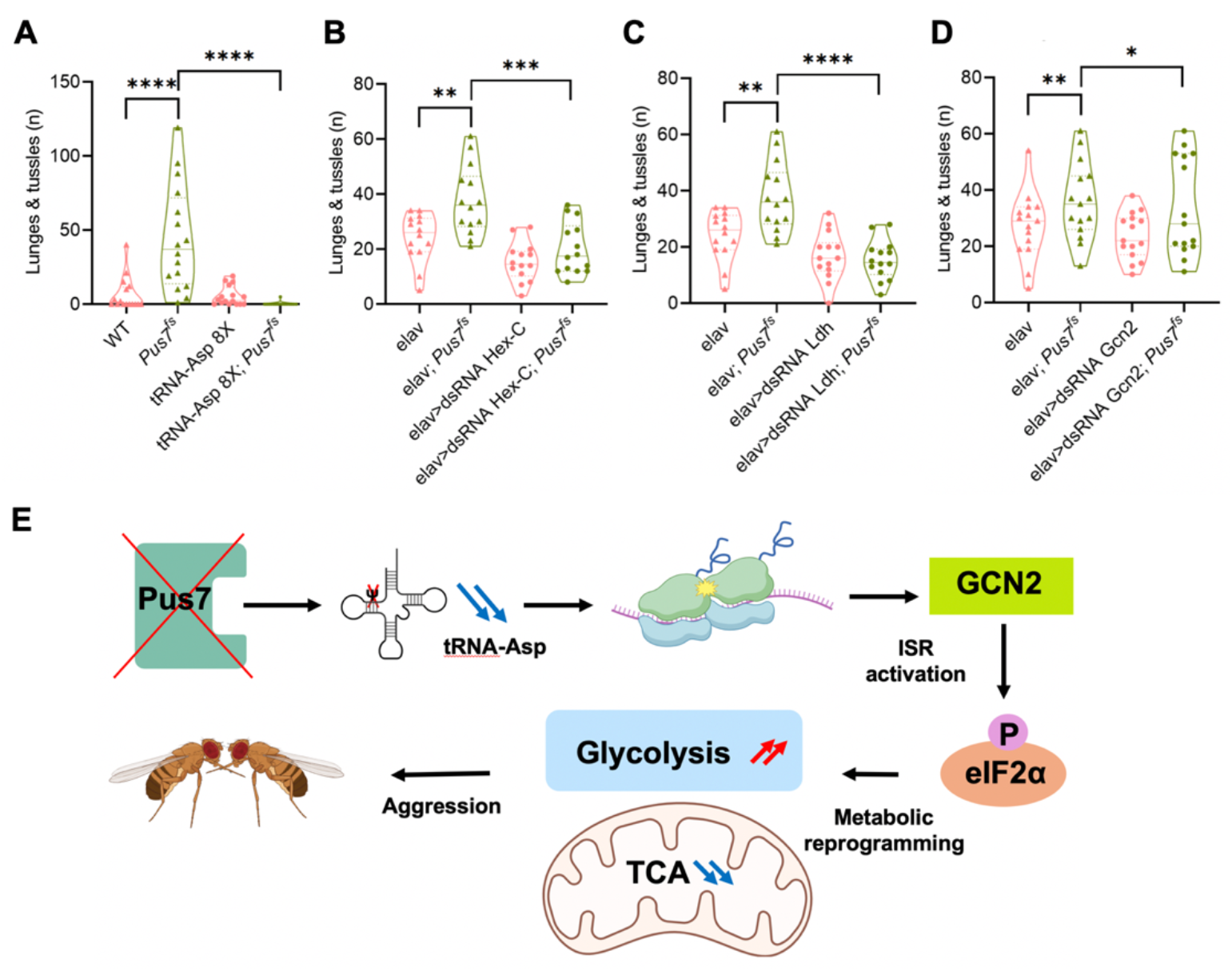
tRNA-Asp, glycolysis and ISR affect aggressiveness in *Pus7*^*fs*^. **flies**. **A**. Violin plot showing the aggression levels (number of lunges and tussles) in WT (n = 15), *Pus7*^*fs*^ (n = 15), tRNA-Asp-GTC 8X (n = 15) and tRNA-Asp 8X; *Pus7*^*fs*^ (n = 15) flies (Mann-Whitney test, P value < 0.0001****, P value > 0.05 ns). **B**. Violin plot showing the aggression levels (number of lunges and tussles) in elav (n = 15), elav; *Pus7*^*fs*^ (n = 15), elav > dsRNA Ldh (n = 15) and elav>dsRNA Ldh; *Pus7*^*fs*^ (n = 15) (Mann-Whitney test, P value < 0.0001 ****, P value < 0.005 **, P value > 0.05 ns). **C**. Violin plot showing the aggression levels (number of lunges and tussles) in elav (n = 15), elav; *Pus7*^*fs*^ (n = 15), elav > dsRNA Hex-C (n = 15) and elav>dsRNA Hex-C; *Pus7*^*fs*^ (n = 15) (Mann-Whitney test, P value < 0.001 ***, P value < 0.005 **, P value > 0.05 ns) (right). **D**. Violin plot showing the aggression levels (number of lunges and tussles) in elav (n = 15), elav; *Pus7*^*fs*^ (n = 15), elav > dsRNA Gcn2 (n = 15) and elav>dsRNA Gcn2; *Pus7*^*fs*^ (n = 15) (Mann-Whitney test, P value < 0.005 **, P value > 0.05 ns).

### Dampening the glycolytic pathway in *Pus7* mutants rescues aggressiveness

To test whether the increased aggression level of *Pus7*^*fs*^ flies could be a direct consequence of the metabolic reprogramming, we attempted to rescue the aggression phenotype by supplementing the food with a glycolytic inhibitor, 2-deoxy-glucose (2-DG). Notably, isolating *Pus7*^*fs*^ males in food supplemented with 100 mM 2-DG was sufficient to ameliorate the aggression levels, indicating that the increase in glycolysis contributed to fly aggressiveness (**Fig. S4A**). By contrast, downregulating OXPHOS by supplementing the food with inhibitory drugs did not increase aggression levels in wild type flies (**Fig. S4B**). We concluded that the causative mechanism was more complex than a simple dysregulation glycolysis:OXPHOS ratios.

Since the drugs act systemically in the organism, thereby likely generating multiple indirect effects, we next attempted to inhibit glycolysis in a more targeted manner. In the *Pus7* mutant background and specifically in neurons, we knocked down the enzyme hexokinase that converts glucose into glucose 6-phosphate. As observed with 2-DG, this was sufficient to restore aggression to the wild type level (**Fig. 4B**). A similar result was obtained when lactate dehydrogenase was inhibited (**Fig. 4C**), further indicating that the increased level of glycolysis specifically in neurons leads to aggressiveness in *Pus7* mutant flies.

Since reducing glycolytic activity was sufficient to alleviate aggression levels, we hypothesized that depriving the mutant flies of sugar might also have a beneficial effect. To test this, we prepared sugar-free food and monitored the aggression levels of *Pus7*^*fs*^ males. As predicted, this intervention reduced aggression in both WT and mutant flies, suggesting that glycolysis driven by dietary sugar plays an important role in this behavior (**Fig. S4C**).

### Inhibiting the integrated stress response restores normal aggression levels

The analysis of up- and down-regulated proteins did not reveal a particular enrichment of Asp codons in their coding sequences, suggesting that many protein abundance changes reflected secondary effects, rather than direct action of Pus7 on Asp decoding (**Fig. S5**). Ribosome stalling and collisions can lead to the activation of stress signaling pathways, notably the ISR, which can result in profound changes to gene expression [29]. The main sensing mechanism in this pathway involves Gcn1 and the eIF2α kinase Gcn2 [30]. Given the slow decoding at Asp codons, as well as the signal of upstream ribosome queueing (**Fig. 2E**), we hypothesized that this pathway may play a role in the phenotypic outcomes, notably on aggressiveness. To test this hypothesis, we fed the flies with ISRIB, an inhibitor of the ISR pathway.

Remarkably, we found that this treatment was sufficient to substantially decrease aggressiveness in *Pus7*^*fs*^ flies (**Fig. S4D**). Likewise, we found that downregulating Gcn2 in the neurons of *Pus7*^*fs*^ flies was sufficient to reduce aggressiveness (**Fig. 4D**). In conclusion, these experiments indicate that activation of the ISR pathway plays a key role in promoting aggressiveness in *Pus7*^*fs*^ flies.

## Discussion

tRNA molecules undergo extensive modifications mediated by tRNA modification enzymes, which are critical for protein synthesis. These enzymes regulate tRNA structure, stability, and the decoding of genetic information on mRNA and their alterations can result in diseases that have been collectively termed tRNA modopathies. These disorders often manifest as brain or kidney dysfunctions, mitochondrial diseases, or cancer. Despite their significance, the molecular mechanisms linking aberrant tRNA modifications to human diseases still remain poorly understood. In this study, we demonstrate that a change in glucose metabolism, stemming from the loss of a specific tRNA modification, plays a causative role in driving complex behavioral changes, such as increased aggressiveness (**Fig. 4E**). This metabolic shift likely arises from the activation of the integrated stress response, triggered by translational impairment at specific codons. Previous associations between other stress related responses and alterations in carbon and lipid metabolism have already been reported [31], [32]. Notably, the unfolded protein response (UPR), activated during endoplasmic reticulum (ER) stress, has been shown to drive the activation of the transcription factor ATF4. This activation leads to changes in the expression of genes involved in carbon metabolism, which include the *Ldh* gene. Interestingly, we observe that *Ldh* transcript levels increase more than threefold in *Pus7* mutant flies, suggesting that ATF4 activation may be the common mediator in these pathways.

Since ISR is frequently activated following disruptions in tRNA modifications, we believe it is worthwhile exploring more systematically its role in tRNA modopathies. This phenomenon may be extended beyond alterations of tRNA modifications. For instance, recent research has also identified ISR activation as a key factor in Charcot-Marie-Tooth disease, which arises from dominant mutations in tRNA synthetase genes [33], [34]. Chronic ISR is also observed in other diseases, for instance upon loss of the Integrator complex that is an essential regulator of transcription whose alteration is associated with many human diseases including cancer and neurodevelopmental disorders [35]. Investigating whether ISR activation leads to metabolic reprogramming that could play a causative role in the progression of these diseases certainly warrants further investigation.

A link between aggressive behavior and metabolism was not previously reported in *Drosophila melanogaster*; however, a similar association has been identified in bees [36]. Here, we demonstrate that increased glycolytic activity directly contributes to aggressiveness. The observation that PUS7 human patients also exhibit increased aggression suggests a conserved mechanism across species, although alterations in the metabolism need yet to be determined. Growing evidence highlights the connections between metabolic changes and human neuronal disorders. For instance, alterations to glucose metabolism have been implicated in Alzheimer’s disease [36], [37]. Additionally, mitochondrial dysfunction, a hallmark of metabolic imbalances, is commonly observed in psychiatric conditions and neurodegenerative diseases [38], [39]. These findings emphasize the potential for metabolic reprogramming to influence neuronal function and behavior. In the case of PUS7 loss-of-function, several key questions remain unresolved. Specifically, the identity of the neuronal populations involved in aggressiveness needs to be determined. Furthermore, the exact mechanism by which the ISR couples to the metabolic switch, the potential role of other PUS7 targets such as mRNA, the consequences of Pus7 mutation in other neuronal phenotypes, and the identification of metabolite(s) potentially driving aggressiveness, all warrant further investigation.

Altogether, the data presented in this study uncover a novel mechanism linking the loss of tRNA modification to behavioral alterations and suggest that inhibiting the ISR pathway and/or reducing the glycolytic activity could represent potential therapeutic strategies for PUS7 patients.

## Materials and Methods

### Drosophila stocks and genetics

*D. melanogaster* reared at 25°C and 60% humidity in a 12 h light/dark cycle. *Drosophila melanogaster* Canton-S with mutant alleles for *Pus7* (*CG6745*) were described previously [16]. Other fly stocks used are *tub*-GAL4, *elavC155*-GAL4 (BL #458), dsRNA Ldh (BL #33640), dsRNA Hex-C (BL #57404), dsRNA Gcn2 (BL #35355) and dsRNA Znf598 (BL #67278). The fly lines for overexpression of tRNA-Asp and tRNA-Lys were generated by FlyORF (Zürich, CH). The U6.2B plasmid containing four copies of tRNA-Asp-GTC-1-5 or four copies of tRNA-Lys-CTT-1-1 was integrated at the attP40 site using the phiC31 system. Fly stocks for behavior experiments were isogenized by crossing to Canton-S at least five times.

### Cloning

The plasmids used for overexpression of Pus7 were constructed by cloning the cDNA of *Pus7* in the Gateway-based vector with N-terminal 3xFLAG–6xMyc tag (pPFMW). The catalytic dead version of this plasmid was generated by introducing a C>G point mutation at position 822 in the CDS of Pus7 changing the catalytic aspartate at position 274 to glutamate. The plasmids for overexpression of tRNA-Asp and tRNA-Lys were generated by cloning two fragments of 393 bp encompassing tRNA-Asp-GTC-1-5 into BbsI digested U6.2 and U6.2B using Gibson assembly or by cloning two fragments of 422 bp encompassing tRNA-Lys-CTT-1-1 into BbsI digested U6.2 and U6.2B using Gibson assembly. Then, the two copies from U6.2 were integrated into U6.2B using EcoRI and NotI to gain the final vector with four copies of tRNA-Asp-GTC-1-5 or to gain the final vector with four copies of tRNA-Lys-CTT-1-1.

### RNA isolation

Total RNA was isolated from flies using TRIzol reagent (Invitrogen) according to manufacturer instructions.

### Northern blot

Total RNA for northern blotting was isolated from flies as described above. Northern blotting was performed essentially as described in [40] with the following adaptations: Fresh 12% urea-polyacrylamide gels were generated using the ROTIPHORESE DNA sequencing system (A431.1, Roth) and polymerized for at least 30 min at RT. Gels were pre-run for at least 30 min in 0.5x tris-borate buffer without EDTA at 90 V. 12 μg of total RNA were prepared in 2x RNA gel loading dye (R0641, Thermo Fisher). 10 fmol of a sense DNA oligo were loaded as a labeling control. RNA samples were denatured at 75°C for 3 min and kept on ice until loading. Electrophoresis was performed at 90 V until the loading dyes visibly separated and finished at 180 V until the bromophenol blue reached the bottom of the gel. Gels were stained in 1x TBE buffer with 1:10000 SYBR-Gold (S11494, Thermo Fisher) for 5 min while shaking at RT. Wet transfer to positively charged nylon membranes (11209299001, Roche) was performed in 0.5x TBE in Bio-Rad mini protean 3 chambers for 90 min at 200 mA. Membranes were dried on whatman paper for at least 5 min before UV crosslinking. Membranes were prehybridized for at least 60 min at 42°C in a screw cap flask in a heated oven in 10 ml hybridization buffer (5x SSC (S6639, Merck), 25 mM Na2HPO4 pH 7.2, 0.5% SDS, 1.25x Denhardts solution (750018, Thermo Fisher). DNA probes were labeled with γ-32P ATP (NEG502A250UC. Perkin Elmer) using T4 PNK (EK0031, Thermo Fisher). Excess ATP was removed using Microspin G-25 columns (GE27-5325-01, Sigma). Hybridization was performed over night at 42°C. Membranes were washed once with Buffer A (3x SSC, 5% SDS) and Buffer B (1x SSC, 1% SDS) for at least 15 min at 42°C, respectively. Finally, membranes were dried on whatman paper, enclosed in plastic wrap and exposed to a phosphor-imaging screen. Signal quantification was performed using ImageJ version 1.53i. Oligonucleotides used for tRNA detection are tRNA31-Asp 5’ AS (CTAACCACTATACTATCGAGGA) tRNA31-Asp 5’ S (TCCTCGATAGTATAGTGGTTAG) and for 5S rRNA are 5S rRNA AS (CCGACCCTGCTTAGCTTCC) and 5S rRNA S (GGAAGCTAAGCAGGGTCGG).

### In vitro tRNA stability assay

*In vitro* transcribed fly tRNAs were prepared as previously described [41]and the stability assays were essentially performed as in [6]. In short, 40 μL of 200nM tRNA was mixed with 200x diluted Quant-iT RiboGreen™ RNA fluorescent probe (R11491 ThermoFisher Scientific) in total 50 μL of the assay buffer (20 mM HEPES, pH 7.5, 150 mM NaCl, 1 mM MgCl_2_, 5 mM DTT). After 10 min of incubation in the dark, 10 μL of the assay mix was loaded into each of the 4 capillaries (NanoTemper Technologies GmbH, cat. no AN-041001). The capillaries were sealed with oil-based liquid rubber (NanoTemper TechnologiesGmbH, cat. no PR-P001) and loaded into the Andromeda system from NanoTemper Technologies. Data were collected with AN.Control Software v1.1 (NanoTemper TechnologiesGmbH, cat. no AN-020001) applying the following settings: total signal intensity at 510 nm adjusted to maximum 2000 counts; the temperature increase of 1ºC/min and fluorescence signal collection in the 25-95 ºC temperature range. The raw data were processed with Origin Pro 2023 (OriginLabs). The signal processing involved linear interpolation to obtain equal size of the temperature vector, Min-Max normalization of the fluorescence signal, and first derivative determination with baseline correction using PeakAnalyzer from OriginLabs.

### Aggression assay

Aggression assays were performed as described in [16], however, only strong aggression events (lunges and tussling) were quantified. Since tussling involved two flies being aggressive, each event was scored twice.

Statistical analysis was performed using GraphPad Prism v8.0.2. For evaluation of aggression, Mann-Whitney U test was used for comparison of two genotypes.

### Psi-seq

To quantify tRNA-Asp levels in both human patient and control samples, as well as in *Drosophila melanogaster* wild type (WT) and *Pus7*^*fs*^ mutants, previously published datasets were used [16]. In addition, Ψ-seq was performed on total RNA isolated from Drosophila heads, with four biological replicates per *Pus3* knockout genotype. Each replicate consisted of 200 ng of total RNA. Ψ-seq was carried out following the protocol described earlier [42].

For tRNA quantification, input (non-CMC-treated) samples were analyzed using the mim-tRNAseq pipeline [43], which enables accurate alignment of highly similar tRNA transcripts. For human samples, the *H. sapiens* hg38 reference genome was used with a cluster identity threshold of 0.97. For fly samples, the *D. melanogaster* dm6 reference genome was used with a cluster ID threshold of 0.9. Read counts generated by the mim-tRNAseq pipeline were further processed to calculate Trimmed Mean of M-values (TMM)-normalized tRNA transcript abundance. Only tRNA transcripts with read counts greater than 10 were included in the final analysis.

### Microscale thermophoresis

Microscale thermophoresis experiments were performed as described in [27]. The total RNA samples were generated from whole flies of mixed sex and age. Oligonucleotides used for tRNA detection are chr3R.trna25Gly TCC (GCGTCGGTGGTGTAATGGTTAGCATAGTTGCCTTCCAAGCAGTTGACCCGGGTTCGATTCCCGGCCGACGCA) and chr2L.trna31Asp GTC (TCCTCGATAGTATAGTGGTTAGTATCCCCGCCTGTCACGCGGGAGACCGGGGTTCAATTCCCCGTCGGGGAG).

### Oroboros assay

Mitochondrial respiration in Drosophila heads was assessed by high resolution respirometry using Oroboros O2k (OROBOROS, Innsbruck, Austria) at 25 **°**C as previously in [44]. Briefly, 15 fly heads from 3-4 day old females, mated were rapidly collected under a microscope, homogenized in MiR05 respiration buffer (20 mM HEPES, 110 mM sucrose, 10 mM KH2PO4, 20 mM taurine, 60 mM lactobionic acid, 3 mM MgCl2, 0.5 mM EGTA, pH 7.1, 1 mg/ml fatty acidfree BSA) using a pestle. The lysate was loaded into an Oroboros O2k chamber with subsequent injections of the following substrates and specific inhibitors: 1) 2.5 mM pyruvate and 1 mM malate (Leak), followed by 2.5 mM ADP to determine complex I-driven phosphorylating respiration (CI OXPHOS). 2) 5 mM succinate to determine the phosphorylating respiration driven by simultaneous activation of complex I and II (CI+CII OXPHOS). 3) Titrating concentrations of the mitochondrial uncoupler CCCP (Carbonyl cyanide 3-chlorophenylhydrazone) to reach the maximal, uncoupled respiration (CI+II electron transfer system, ETS). 4) 200 nM rotenone to fully inhibit complex I-driven respiration and measure complex II-driven uncoupled respiration (CII electron transfer system, CII ETS). 5) 0.5 μM Antimycin A to block mitochondrial respiration at the level of complex III. Residual oxygen consumption (ROX) was always negligible. 6) 2 mM ascorbate, 0.5 mM TMPD (N,N,N’,N,- Tetramethyl-p phenylenediamine) to measure cytochrome c oxidase (CIV or COX)-driven respiration. 7) 300 μM Sodium Azide to specifically block cytochrome c oxidase activity and measure residual background oxygen consumption caused by chemical interaction between ascorbate and TMPD. For the analysis ROX was subtracted from all the substrate-specific oxygen consumption rates (Leak, CI OXPHOS, CI-CII OXPHOS) while the maximal oxygen consumption rate of complex IV was corrected by subtracting the complex IV background. O2 consumption rates were normalized over total protein content assessed by Pierce BCA Protein Assay Kit (ThermoFisher).

### Ribosome profiling and bioinformatic analysis

RNA samples for ribosome profiling were prepared as described in [45] with the adaption that customized siTOOLs depletion pools for Drosophila rRNA were used. Libraries were generated from approximately 80 flash frozen heads of 3-4 day old mated female flies in technical triplicates. The codon occupancy analysis for P and A sites was performed as described in [46]. Putative ribosome pausing sites were identified as described in [47].

### Proteomics and bioinformatic analysis

### Sample preparation and protein digestion

Five replicates of 20 heads per replicate and genotype of 3-4-day old mated females were prepared by flash freezing whole flies in eppendorf tubes in liquid nitrogen and separating heads by repeatedly smashing the frozen tube on to a table. Heads separated from fly bodies were collected with a brush and immediately frozen in liquid nitrogen. Drosophila heads were crushed manually with a pestel in 1.5 ml tubes in 120 ul of miST lysis buffer (1% sodium deoxycholate, 100 mM Tris pH 8.6, 10 mM DTT). The suspension was heated at 95°C for 5 min, cooled down and sonicated with a tip sonicator for 20 s on ice, resulting in homogeneous suspensions. Consistency of protein extraction was controlled qualitatively by 1D-SDS-PAGE (data not shown). Samples were digested following a modified version of the iST method (named miST method) [48]. Based on tryptophane fluorescence quantification [49], 100 μg of proteins at 2 μg/μl in miST lysis buffer were transferred to new tubes. Samples were heated 5 min at 95°C, diluted 1:1 (v:v) with water containing 4 mM MgCl2 and benzonase (Merck #70746, 100x dil of stock = 250 Units/μl), and incubated for 15 minutes at RT to digest nucleic acids. Reduced disulfides were alkylated by adding 1/4 vol. of 160 mM chloroacetamide (32 mM final) and incubating for 45 min at RT in the dark. Samples were adjusted to 3 mM EDTA and digested with 1 μg Trypsin/LysC mix (Promega) #V5073) for 1 h at 37°C, followed by a second 1 h digestion with an additional 1 μg of proteases. To remove sodium deoxycholate, two sample volumes of isopropanol containing 1% TFA were added to the digests, and the samples were desalted on a strong cation exchange (SCX) plate (Oasis MCX; Waters Corp., Milford, MA) by centrifugation. After washing with isopropanol/1%TFA, peptides were eluted in 200 ul of 80% MeCN, 19% water, 1% (v/v) ammonia, and dried by centrifugal evaporation.

### Peptide fractionation

Aliquots of samples were pooled and separated into 6 fractions by off-line basic reversed-phase (bRP) using the Pierce High pH Reversed-Phase Peptide Fractionation Kit (Thermo Fisher Scientific). The fractions were collected in 7.5, 10, 12.5, 15, 17.5 and 50% acetonitrile in 0.1% triethylamine (pH 10). Dried bRP fractions were redissolved in 50 μl 2% acetonitrile with 0.5% TFA, and 5 μl were injected for LC-MS/MS analyses.

### Liquid Chromatography-Mass Spectrometry analyses

LC-MS/MS analyses were carried out on a TIMS-TOF Pro (Bruker, Bremen, Germany) mass spectrometer interfaced through a nanospray ion source (“captive spray”) to an Ultimate 3000 RSLCnano HPLC system (Dionex). Peptides were separated on a reversed-phase custom packed 45 cm C18 column (75 μm ID, 100Å, Reprosil Pur 1.9 um particles, Dr. Maisch, Germany) at a flow rate of 0.250 μl/min with a 2-27% acetonitrile gradient in 93 min followed by a ramp to 45% in 15 min and to 90% in 5 min (total method time: 140 min, all solvents contained 0.1% formic acid). Identical LC gradients were used for DDA and DIA measurements. For creation of the spectral library, data dependent acquisitions (DDA) were carried out on the 6 bRP fractions sample pool using a standard TIMS PASEF method [50] with ion accumulation for 100 ms for each survey MS1 scan and the TIMS-coupled MS2 scans. Duty cycle was kept at 100%. Up to 10 precursors were targeted per TIMS scan. Precursor isolation was done with a 2 Th or 3 Th windows below or above m/z 800, respectively. The minimum threshold intensity for precursor selection was 2500. If the inclusion list allowed it, precursors were targeted more than one time to reach a minimum target total intensity of 20.000. Collision energy was ramped linearly based uniquely on the 1/k0 values from 20 (at 1/k0=0.6) to 59 eV (at 1/k0=1.6). Total duration of a scan cycle including one survey and 10 MS2 TIMS scans was 1.16 s. Precursors could be targeted again in subsequent cycles if their signal increased by a factor 4.0 or more. After selection in one cycle, precursors were excluded from further selection for 60 s. Mass resolution in all MS measurements was approximately 35.000. The data-independent acquisition (DIA) used mostly the same instrument parameters as the DDA method and was as reported previously [51]. Per cycle, the mass range 400-1200 m/z was covered by a total of 32 windows, each 25 Th wide and a 1/k0 range of 0.3. Collision energy and resolution settings were the same as in the DDA method. Two windows were acquired per TIMS scan (100 ms) so that the total cycle time was 1.7 s

### Data processing

Raw Bruker MS data were processed directly with Spectronaut 15.6 (Biognosys, Schlieren, Switzerland). A library was constructed from the DDA bRP fraction data by searching the reference Drosophila melanogaster proteome (www.uniprot.org) database of August 23rd, 2020 (22.039 sequences). For identification, peptides of 7-52 AA length were considered, cleaved with trypsin/P specificity and a maximum of 2 missed cleavages. Carbamidomethylation of cysteine (fixed), methionine oxidation and N-terminal protein acetylation (variable) were the modifications applied. Mass calibration was dynamic and based on a first database search. The Pulsar engine was used for peptide identification. Protein inference was performed with the IDPicker algorithm. Spectra, peptide and protein identifications were all filtered at 1% FDR against a decoy database. Specific filtering for library construction removed fragments corresponding to less than 3 AA and fragments outside the 300-1800 m/z range. Also, only fragments with a minimum base peak intensity of 5% were kept. Precursors with less than 3 fragments were also eliminated and only the best 6 fragments were kept per precursor. No filtering was done on the basis of charge state and a maximum of 2 missed cleavages was allowed. Shared (non proteotypic) peptides were kept. The library created contained 79.159 precursors mapping to 53.911 peptides, of which 27.089 were proteotypic. These corresponded to 6.459 protein groups (9.690 proteins). Of these, 941 were single hits (one peptide precursor). In total 465.866 fragments were used for quantitation. Peptide-centric analysis of DIA data was done with Spectronaut 15.6 using the library described above. Single hits proteins (defined as matched by one stripped sequence only) were kept in the Spectronaut analysis. Peptide quantitation was based on XIC area, for which a minimum of 1 and a maximum of 3 (the 3 best) precursors were considered for each peptide, from which the median value was selected. Quantities for protein groups were derived from inter-run peptide ratios based on MaxLFQ algorithm [52]. Global normalization of runs/samples was done based on the median of peptides.

### Data analysis

All subsequent analyses were done with the Perseus software package (version 1.6.15.0) [53]. Contaminant proteins were removed, and intensity values determined by Spectronaut were log2-transformed. After assignment to groups, only proteins quantified in at least 4 samples in at least one group were kept (6.130 protein groups). After imputation of missing values (based on normal distribution using Perseus default parameters), t-tests were carried out among all conditions, with permutation-based FDR correction for multiple testing (q-value <0.05). 838 proteins passed the test with these conditions, including Pus7. The difference of means obtained from the tests were used for 1D enrichment analysis on associated GO/KEGG annotations as described [54]. The enrichment analysis was also FDR-filtered (Benjamini-Hochberg, q-val <0.02).

### Gene ontology (GO) analysis of proteomics data

GO term analysis was performed with GO Enrichment analysis tool [55], [56], [57].

### Multiple pathway targeted metabolomics Metabolite extraction

Six replicates of 50 Fly heads of 3-4 day old mated females were pre-extracted and homogenized by the addition of 150 μL of MeOH:H2O (4:1), in the Cryolys Precellys 24 sample Homogenizer (2 × 20 seconds at 10000 rpm, Bertin Technologies, Rockville, MD, US) with ceramic beads. The bead beater was air-cooled down at a flow rate of 110 L/min at 6 bar. Homogenized extracts were centrifuged for 15 minutes at 4000 g at 4°C (Hermle, Gosheim, Germany). The resulting supernatant was collected and injected into the LC-MS system.

### LC-MS/MS analysis

Sample extracts were analyzed by Hydrophilic Interaction Liquid Chromatography coupled to tandem mass spectrometry (HILIC - MS/MS) in both positive and negative ionization modes using a 6495 triple quadrupole system (QqQ) interfaced with 1290 UHPLC system (Agilent Technologies) [58], [59]. The extracts were analyzed by hydrophilic interaction chromatography coupled to tandem mass spectrometer (6496 iFunnel Agilent Tehcnologies) using multiple reaction monitoring - MRM approach in both, positive and negative ionization modes, to maximize the polar metabolome coverage. Detailed analytical conditions have been previously described in [58], [59].

### Data processing and quality assessment

Raw LC-MS/MS data were processed using the Mass Hunter Quantitative analysis software (Agilent Technologies). The peak areas (or extracted ion chromatograms (EICs) for the monitored MRM transitions) were used for relative comparison of metabolite levels between different conditions or groups of samples. Data quality assessment, including signal drift correction, was performed using pooled quality control (QC) samples analyzed periodically throughout the entire batch. Signals with analytical CV<30% were discarded.

### Resource availability

Further information and requests for resources should be directed to the lead contact, Jean-Yves Roignant

## Data and code availability

All data needed to evaluate the conclusions in the paper are present in the paper and/or Supplemental materials. Proteomics data are available in PRIDE (PXD063492). Additional data related to this paper may be requested from the authors.

## Acknowledgments

We thank the Bloomington stock center for fly lines and FlyORF for injections. We thank the members of the Roignant lab and collaborators for helpful discussion. Research in the laboratory of JYR is supported by the University of Lausanne, The Swiss National Science Foundation (310030_197906), by Deutsche Forschungsgemeinschaft Collaborative Research Center RMaP (TRR 319, TP B01). DG acknowledges funding by SNSF individual grant (212423) and National Center for Competence in Research (NCCR) RNA & Disease (205601). Research in the lab of SG was provided by the European Research Council (ERC) under the European Union’s Horizon 2020 research and innovation program grant No 101001394. MH was funded by grants from the Deutsche Forschungsgemeinschaft (DFG, German Research Foundation) project number 439669440 TRR319 RMaP TP C01 and C03.

## Author contributions

MS, MGA and JYR conceived the project. All authors contributed to the experimental design, analysis, and interpretation of results. Experimental contributions were as follows: MS and MGA performed all experiments except the following ones: MW and SS performed the Psi-Seq experiment and analysis. AL and DG performed the Ribo-Seq. AB, JN, ACG and SG carried out the in vitro tRNA stability assay. VM and CB carried out the Oroboros experiment. DJ, MK and MH performed the microscale thermophoresis assay. MQ carried out the proteomics experiment and analysis. AS, RD, ABr and NG performed the bioinformatic analysis. HGA and JI performed the metabolomics assay. LK and SL contributed to the Ribo-Seq data and interpretation. MS, MGA and JYR wrote the paper with input from all the authors.

## Declaration of interest

The authors declare no competing interests

---

**Supplementary figure 1.**
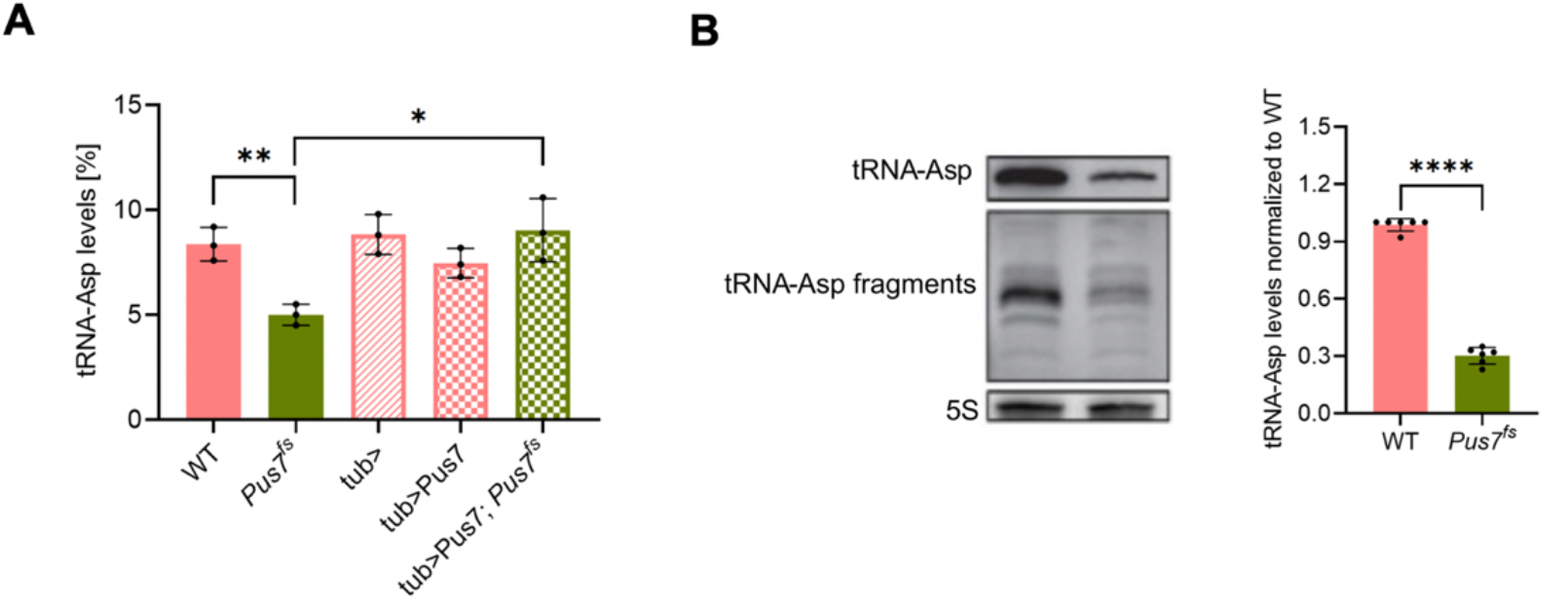
Loss of Pus7 leads to reduced tRNA-Asp levels. **A**. Bar plot showing tRNA-Asp levels (%) measured by microscale thermophoresis in WT (n = 3), *Pus7*^*fs*^ (n = 3), tub> (n = 3), tub>Pus7 (n = 3) and tub>Pus7; *Pus7*^*fs*^ (n = 3) flies. **B**. (left) Northern blot membrane showing tRNA-Asp mature and fragment levels in WT and *Pus7*^*fs*^. (right) Bar plot showing tRNA-Asp levels by relative intensity normalized to WT measured by NB in WT (n = 6) and *Pus7*^*fs*^ (n = 6) (t-test unpaired, value < 0.0001 ****) (top)

**Supplementary figure 2.**
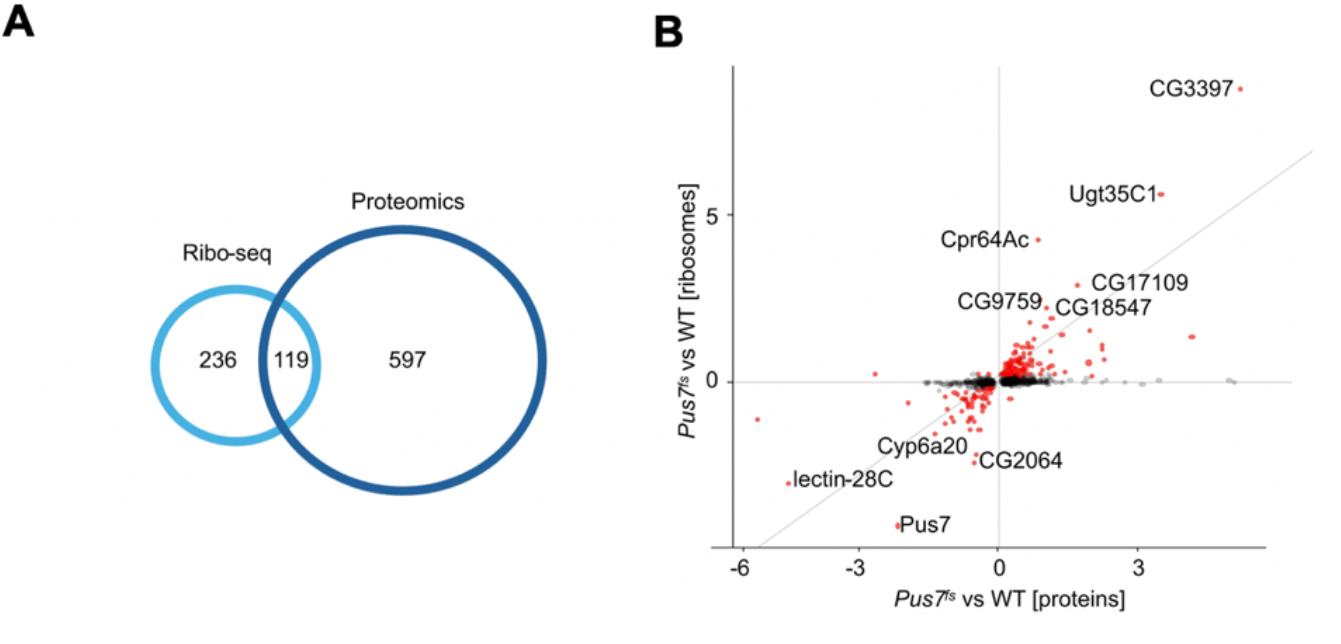
Correlation between Ribo-seq and proteomics. **A**. Overlap of differentially expressed transcripts found in Ribo-Seq and proteins found in protein mass spectrometry in *Pus7*^*fs*^ compared to wild type fly heads. **(B)** Scatter plot showing correlation of differentially expressed transcripts found in Ribo-Seq and proteins in protein mass spectrometry (bottom) in *Pus7*^*fs*^ compared to wild type fly heads.

**Supplementary figure 3.**
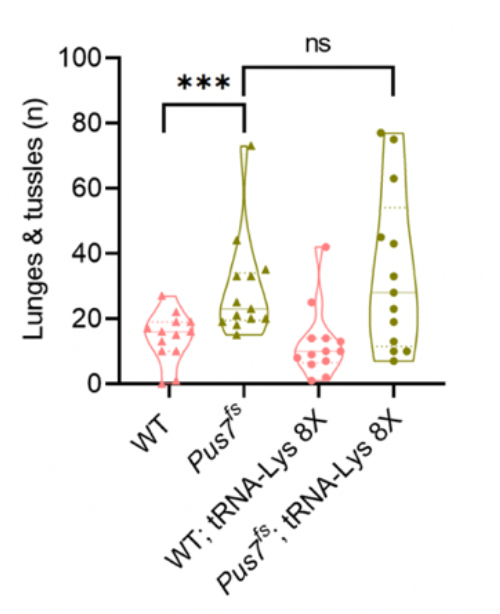
Overexpression of tRNA-Lys does not rescue aggression in *Pus7* mutant flies. Violin plot showing the aggression levels (number of lunges and tussles) in WT (n = 15), *Pus7*^*fs*^ (n= 15), tRNA-Lys 8X (n = 15) and tRNA-Lys 8X; *Pus7*^*fs*^ (n = 15) flies (Mann-Whitney test, P value < 0.001***, P value > 0.05 ns).

**Supplementary figure 4.**
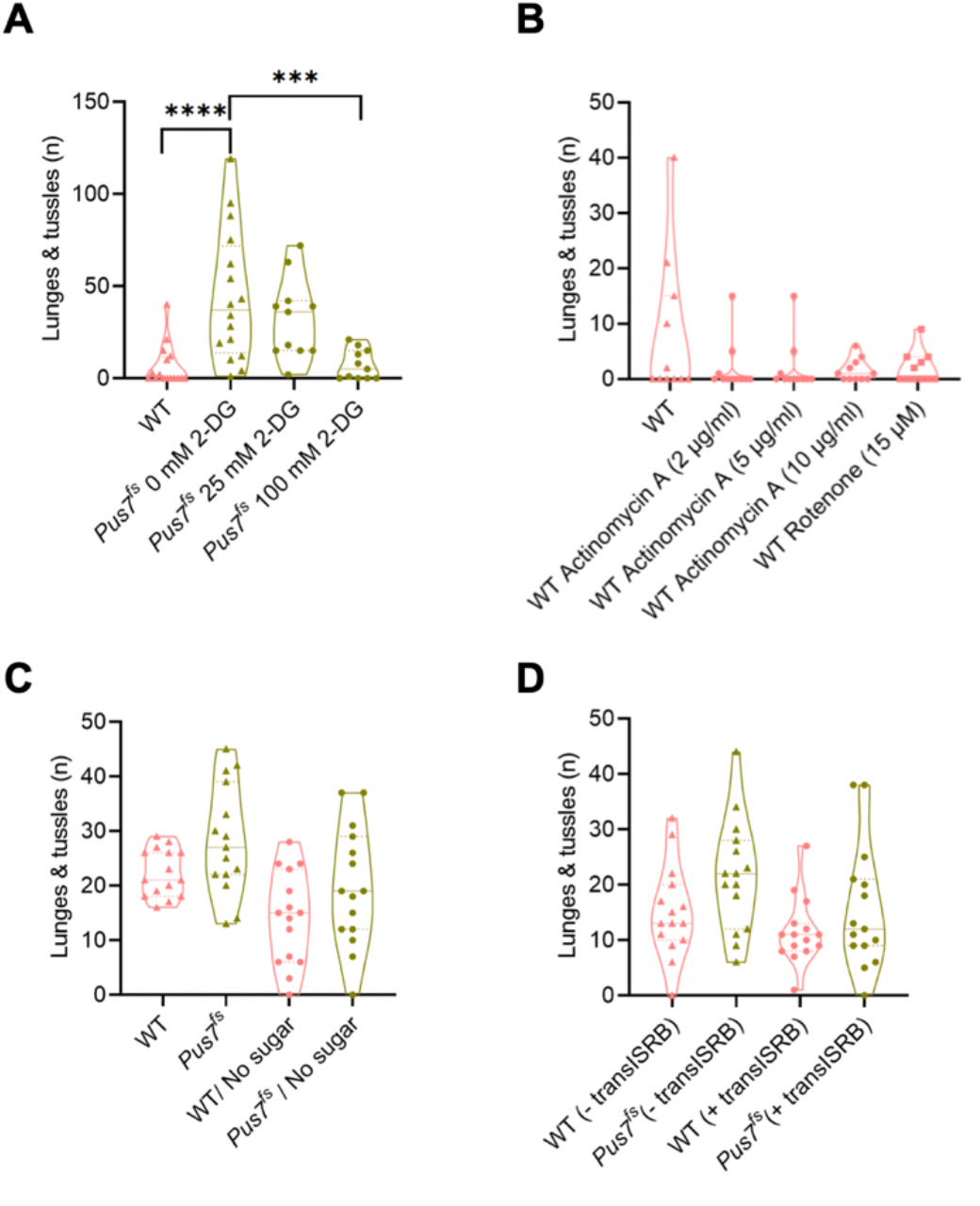
Aggression in *Pus7* mutant results from ISR activation and increased glycolysis. **A**. Violin plot showing the aggression levels (number of lunges and tussles) in WT (n = 15), *Pus7*^*fs*^ (n = 15) exposed to 0 mM of DG in food, *Pus7*^*fs*^ (n = 15) exposed to 25 mM of DG in food (n = 15) and *Pus7*^*fs*^ (n = 15) exposed to 100 mM of DG in food (Mann-Whitney test, P value < 0.001***). **B**. Violin plot showing the aggression levels (number of lunges and tussles) in WT (n = 15), WT (n = 15) exposed to 15 μM of rotenone in food, WT (n = 15) exposed to (2 μg/ml) of actinomycin A in food (n = 15), WT (n = 15) exposed to (5 μg/ml) of actinomycin A of in food and WT (n = 15) exposed to (10 μg/ml) of actinomycin A of in food (Mann-Whitney test, P value > 0.05 ns). **C**. Violin plot showing the aggression levels (number of lunges and tussles) in WT (n = 15) exposed to standard food recipe, *Pus7*^*fs*^ (n = 15) exposed to standard food recipe, WT (n = 15) exposed to food without glucose and *Pus7*^*fs*^ (n = 15) exposed to food without glucose (Mann-Whitney test, P value > 0.05 ns). **D**. Violin plot showing the aggression levels (number of lunges and tussles) in WT (n = 15), *Pus7*^*fs*^ (n = 15), *Pus7*^*fs*^ (n = 15) exposed to 500 nM of ISRB in food (n = 15) and *Pus7*^*fs*^ (n = 15) exposed to 500 nM of ISRB in food (Mann-Whitney test, P value > 0.05 ns).

**Supplementary figure 5.**
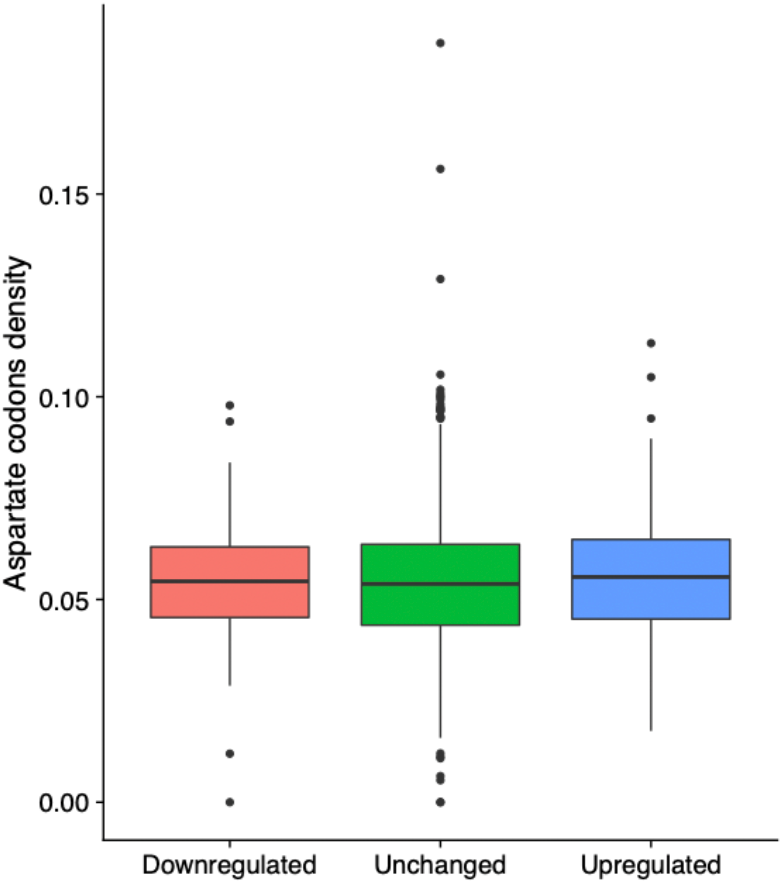
Aspartate codons are not enriched in differentially regulated proteins. Box plot showing the aspartate codon density in proteins found to be upregulated, downregulated or expression not changed in *Pus7*^*fs*^ compared to wild type fly heads.

## Notes

### Competing Interest Statement

The authors have declared no competing interest.

## References

[1] A. Bednářová et al., ‘Lost in Translation: Defects in Transfer RNA Modifications and Neurological Disorders’, Front Mol Neurosci, vol. 10, p. 135, May 2017, doi: 10.3389/fnmol.2017.00135.

[2] M. T. Angelova et al., ‘The Emerging Field of Epitranscriptomics in Neurodevelopmental and Neuronal Disorders’, Front Bioeng Biotechnol, vol. 6, p. 46, 2018, doi: 10.3389/fbioe.2018.00046.

[3] T.-Y. Lin, R. Mehta, and S. Glatt, ‘Pseudouridines in RNAs: switching atoms means shifting paradigms’, FEBS Lett, vol. 595, no. 18, pp. 2310–2322, Sep. 2021, doi: 10.1002/1873-3468.14188.

[4] M. I. Newby and N. L. Greenbaum, ‘Sculpting of the spliceosomal branch site recognition motif by a conserved pseudouridine’, Nat Struct Mol Biol, vol. 9, no. 12, pp. 958–965, Dec. 2002, doi: 10.1038/nsb873.

[5] M. Sumita, J.-P. Desaulniers, Y.-C. Chang, H. M.-P. Chui, L. Clos, and C. S. Chow, ‘Effects of nucleotide substitution and modification on the stability and structure of helix 69 from 28S rRNA’, RNA, vol. 11, no. 9, pp. 1420–1429, Sep. 2005, doi: 10.1261/rna.2320605.

[6] A. D. Biela et al., ‘Determining the effects of pseudouridine incorporation on human tRNAs’, EMBO J, Apr. 2025, doi: 10.1038/s44318-025-00443-y.

[7] T. Hamma and A.R. Ferré-D’Amaré, ‘Pseudouridine synthases’, Chem Biol, vol. 13, no. 11, pp. 1125–1135, Nov. 2006, doi: 10.1016/j.chembiol.2006.09.009.

[8] A. C. Rintala-Dempsey and U. Kothe, ‘Eukaryotic stand-alone pseudouridine synthases – RNA modifying enzymes and emerging regulators of gene expression?’, RNA Biol, vol. 14, no. 9, pp. 1185–1196, Jan. 2017, doi: 10.1080/15476286.2016.1276150.

[9] L. Huang, M. Pookanjanatavip, X. Gu, and D. V. Santi, ‘A conserved aspartate of tRNA pseudouridine synthase is essential for activity and a probable nucleophilic catalyst’, Biochemistry, vol. 37, no. 1, pp. 344–351, Jan. 1998, doi: 10.1021/bi971874+.

[10] M. Guillen-Angel and J.-Y. Roignant, ‘Exploring pseudouridylation: dysregulation in disease and therapeutic potential’, Curr Opin Genet Dev, vol. 87, p. 102210, Aug. 2024, doi: 10.1016/j.gde.2024.102210.

[11] Y. Bykhovskaya, K. Casas, E. Mengesha, A. Inbal, and N. Fischel-Ghodsian, ‘Missense mutation in pseudouridine synthase 1 (PUS1) causes mitochondrial myopathy and sideroblastic anemia (MLASA)’, Am J Hum Genet, vol. 74, no. 6, pp. 1303–1308, Jun. 2004, doi: 10.1086/421530.

[12] J. Woods and S. Cederbaum, ‘Myopathy, lactic acidosis and sideroblastic anemia 1 (MLASA1): A 25-year follow-up’, Mol Genet Metab Rep, vol. 21, p. 100517, Dec. 2019, doi: 10.1016/j.ymgmr.2019.100517.

[13] T.-Y. Lin et al., ‘Destabilization of mutated human PUS3 protein causes intellectual disability’, Human Mutation, vol. 43, no. 12, pp. 2063–2078, 2022, doi: 10.1002/humu.24471.

[14] R. Shaheen et al., ‘A homozygous truncating mutation in PUS3 expands the role of tRNA modification in normal cognition’, Hum Genet, vol. 135, no. 7, pp. 707–713, Jul. 2016, doi: 10.1007/s00439-016-1665-7.

[15] H. Darvish et al., ‘A novel PUS7 mutation causes intellectual disability with autistic and aggressive behaviors’, Neurol Genet, vol. 5, no. 5, p. e356, Sep. 2019, doi: 10.1212/NXG.0000000000000356.

[16] A. P. M. de Brouwer et al., ‘Variants in PUS7 Cause Intellectual Disability with Speech Delay, Microcephaly, Short Stature, and Aggressive Behavior’, The American Journal of Human Genetics, vol. 103, no. 6, pp. 1045–1052, Dec. 2018, doi: 10.1016/j.ajhg.2018.10.026.

[17] S. T. Han et al., ‘PUS7 deficiency in human patients causes profound neurodevelopmental phenotype by dysregulating protein translation’, Mol Genet Metab, vol. 135, no. 3, pp. 221–229, Mar. 2022, doi: 10.1016/j.ymgme.2022.01.103.

[18] A. Muda, L. Malerba, L. Giordano, E. Fazzi, and P. Accorsi, ‘A PUS7 gene pathogenic variant causing self-injurious behavior, sleep disturbances, and developmental delay: A case report’, Am J Med Genet A, vol. 191, no. 7, pp. 1953–1958, Jul. 2023, doi: 10.1002/ajmg.a.63212.

[19] M. I. Naseer, A. A. Abdulkareem, M. M. Jan, A. G. Chaudhary, S. Alharazy, and M. H. AlQahtani, ‘Next generation sequencing reveals novel homozygous frameshift in PUS7 and splice acceptor variants in AASS gene leading to intellectual disability, developmental delay, dysmorphic feature and microcephaly’, Saudi J Biol Sci, vol. 27, no. 11, pp. 3125–3131, Nov. 2020, doi: 10.1016/j.sjbs.2020.09.033.

[20] R. Shaheen et al., ‘PUS7 mutations impair pseudouridylation in humans and cause intellectual disability and microcephaly’, Hum Genet, vol. 138, no. 3, pp. 231–239, Mar. 2019, doi: 10.1007/s00439-019-01980-3.

[21] N. Guzzi et al., ‘Pseudouridylation of tRNA-Derived Fragments Steers Translational Control in Stem Cells’, Cell, vol. 173, no. 5, pp. 1204-1216.e26, May 2018, doi: 10.1016/j.cell.2018.03.008.

[22] Q. Cui et al., ‘Targeting PUS7 suppresses tRNA pseudouridylation and glioblastoma tumorigenesis’, Nat Cancer, vol. 2, no. 9, Art. no. 9, Sep. 2021, doi: 10.1038/s43018-021-00238-0.

[23] G. Zhang et al., ‘Higher expression of pseudouridine synthase 7 promotes non-small cell lung cancer progression and suggests a poor prognosis’, J Cardiothorac Surg, vol. 18, p. 222, Jul. 2023, doi: 10.1186/s13019-023-02332-z.

[24] J. Du, A. Gong, X. Zhao, and G. Wang, ‘Pseudouridylate Synthase 7 Promotes Cell Proliferation and Invasion in Colon Cancer Through Activating PI3K/AKT/mTOR Signaling Pathway’, Dig Dis Sci, vol. 67, no. 4, pp. 1260–1270, Apr. 2022, doi: 10.1007/s10620-021-06936-0.

[25] D. Song et al., ‘HSP90-dependent PUS7 overexpression facilitates the metastasis of colorectal cancer cells by regulating LASP1 abundance’, Journal of Experimental & Clinical Cancer Research, vol. 40, no. 1, p. 170, May 2021, doi: 10.1186/s13046-021-01951-5.

[26] Q. Zhang et al., ‘PUS7 promotes the proliferation of colorectal cancer cells by directly stabilizing SIRT1 to activate the Wnt/β-catenin pathway’, Mol Carcinog, vol. 62, no. 2, pp. 160–173, Feb. 2023, doi: 10.1002/mc.23473.

[27] D. Jacob et al., ‘Absolute Quantification of Noncoding RNA by Microscale Thermophoresis’, Angewandte Chemie International Edition, vol. 58, no. 28, pp. 9565–9569, 2019, doi: 10.1002/anie.201814377.

[28] C. Lorenz, C. E. Lünse, and M. Mörl, ‘tRNA Modifications: Impact on Structure and Thermal Adaptation’, Biomolecules, vol. 7, no. 2, p. 35, Apr. 2017, doi: 10.3390/biom7020035.

[29] S. Meydan and N. R. Guydosh, ‘Disome and Trisome Profiling Reveal Genome-wide Targets of Ribosome Quality Control’, Molecular Cell, vol. 79, no. 4, pp. 588-602.e6, Aug. 2020, doi: 10.1016/j.molcel.2020.06.010.

[30] L. L. Yan and H. S. Zaher, ‘Ribosome quality control antagonizes the activation of the integrated stress response on colliding ribosomes’, Molecular Cell, vol. 81, no. 3, pp. 614-628.e4, Feb. 2021, doi: 10.1016/j.molcel.2020.11.033.

[31] J. E. Lee, M. Oney, K. Frizzell, N. Phadnis, and J. Hollien, ‘Drosophila melanogaster activating transcription factor 4 regulates glycolysis during endoplasmic reticulum stress’, G3 (Bethesda), vol. 5, no. 4, pp. 667–675, Feb. 2015, doi: 10.1534/g3.115.017269.

[32] S. Sorge et al., ‘ATF4-Induced Warburg Metabolism Drives Over-Proliferation in Drosophila’, Cell Rep, vol. 31, no. 7, p. 107659, May 2020, doi: 10.1016/j.celrep.2020.107659.

[33] A. Zuko et al., ‘tRNA overexpression rescues peripheral neuropathy caused by mutations in tRNA synthetase’, Science, vol. 373, no. 6559, pp. 1161–1166, Sep. 2021, doi: 10.1126/science.abb3356.

[34] S. Mendonsa, N. von Kuegelgen, L. Bujanic, and M. Chekulaeva, ‘Charcot-Marie-Tooth mutation in glycyl-tRNA synthetase stalls ribosomes in a pre-accommodation state and activates integrated stress response’, Nucleic Acids Res, vol. 49, no. 17, pp. 10007–10017, Sep. 2021, doi: 10.1093/nar/gkab730.

[35] A. Baluapuri et al., ‘Integrator loss leads to dsRNA formation that triggers the integrated stress response’, Cell, pp. S0092-8674(25)00343–5, Apr. 2025, doi: 10.1016/j.cell.2025.03.025.

[36] X. Zhang, N. Alshakhshir, and L. Zhao, ‘Glycolytic Metabolism, Brain Resilience, and Alzheimer’s Disease’, Front Neurosci, vol. 15, p. 662242, 2021, doi: 10.3389/fnins.2021.662242.

[37] P. S. Minhas et al., ‘Restoring hippocampal glucose metabolism rescues cognition across Alzheimer’s disease pathologies’, Science, vol. 385, no. 6711, p. eabm6131, Aug. 2024, doi: 10.1126/science.abm6131.

[38] M. M. Klemmensen, S. H. Borrowman, C. Pearce, B. Pyles, and B. Chandra, ‘Mitochondrial dysfunction in neurodegenerative disorders’, Neurotherapeutics, vol. 21, no. 1, p. e00292, Jan. 2024, doi: 10.1016/j.neurot.2023.10.002.

[39] P. Ni, Y. Ma, and S. Chung, ‘Mitochondrial dysfunction in psychiatric disorders’, Schizophr Res, vol. 273, pp. 62–77, Nov. 2024, doi: 10.1016/j.schres.2022.08.027.

[40] D. C. Rio, ‘Northern Blots for Small RNAs and MicroRNAs’, Cold Spring Harb Protoc, vol. 2014, no. 7, p. pdb.prot080838, Jan. 2014, doi: 10.1101/pdb.prot080838.

[41] A. Chramiec-Głąbik, M. Rawski, S. Glatt, and T.-Y. Lin, ‘Electrophoretic Mobility Shift Assay (EMSA) and Microscale Thermophoresis (MST) Methods to Measure Interactions Between tRNAs and Their Modifying Enzymes’, Methods Mol Biol, vol. 2666, pp. 29–53, 2023, doi: 10.1007/978-1-0716-3191-1_3.

[42] S. Schwartz et al., ‘Transcriptome-wide Mapping Reveals Widespread Dynamic-Regulated Pseudouridylation of ncRNA and mRNA’, Cell, vol. 159, no. 1, pp. 148–162, Sep. 2014, doi: 10.1016/j.cell.2014.08.028.

[43] A. Behrens, G. Rodschinka, and D. D. Nedialkova, ‘High-resolution quantitative profiling of tRNA abundance and modification status in eukaryotes by mim-tRNAseq’, Molecular Cell, vol. 81, no. 8, pp. 1802-1815.e7, Apr. 2021, doi: 10.1016/j.molcel.2021.01.028.

[44] A. K. Kanellopoulos et al., ‘Aralar Sequesters GABA into Hyperactive Mitochondria, Causing Social Behavior Deficits’, Cell, vol. 180, no. 6, pp. 1178-1197.e20, Mar. 2020, doi: 10.1016/j.cell.2020.02.044.

[45] G. Snieckute et al., ‘Ribosome stalling is a signal for metabolic regulation by the ribotoxic stress response’, Cell Metabolism, vol. 34, no. 12, pp. 2036-2046.e8, Dec. 2022, doi: 10.1016/j.cmet.2022.10.011.

[46] A. B. Arpat, A. Liechti, M. De Matos, R. Dreos, P. Janich, and D. Gatfield, ‘Transcriptome-wide sites of collided ribosomes reveal principles of translational pausing’, Genome Res, vol. 30, no. 7, pp. 985–999, Jul. 2020, doi: 10.1101/gr.257741.119.

[47] K. C. Stein, F. Morales-Polanco, J. van der Lienden, T. K. Rainbolt, and J. Frydman, ‘Ageing exacerbates ribosome pausing to disrupt cotranslational proteostasis’, Nature, vol. 601, no. 7894, pp. 637–642, Jan. 2022, doi: 10.1038/s41586-021-04295-4.

[48] N. A. Kulak, G. Pichler, I. Paron, N. Nagaraj, and M. Mann, ‘Minimal, encapsulated proteomic-sample processing applied to copy-number estimation in eukaryotic cells’, Nat Methods, vol. 11, no. 3, pp. 319–324, Mar. 2014, doi: 10.1038/nmeth.2834.

[49] J.R. Wiśniewski and F. Z. Gaugaz, ‘Fast and Sensitive Total Protein and Peptide Assays for Proteomic Analysis’, Anal. Chem., vol. 87, no. 8, pp. 4110–4116, Apr. 2015, doi: 10.1021/ac504689z.

[50] F. Meier et al., ‘Online Parallel Accumulation-Serial Fragmentation (PASEF) with a Novel Trapped Ion Mobility Mass Spectrometer’, Mol Cell Proteomics, vol. 17, no. 12, pp. 2534–2545, Dec. 2018, doi: 10.1074/mcp.TIR118.000900.

[51] F. Meier et al., ‘diaPASEF: parallel accumulation–serial fragmentation combined with data-independent acquisition’, Nat Methods, vol. 17, no. 12, pp. 1229–1236, Dec. 2020, doi: 10.1038/s41592-020-00998-0.

[52] J. Cox, M. Y. Hein, C. A. Luber, I. Paron, N. Nagaraj, and M. Mann, ‘Accurate Proteome-wide Label-free Quantification by Delayed Normalization and Maximal Peptide Ratio Extraction, Termed MaxLFQ’, Mol Cell Proteomics, vol. 13, no. 9, pp. 2513–2526, Sep. 2014, doi: 10.1074/mcp.M113.031591.

[53] S. Tyanova et al., ‘The Perseus computational platform for comprehensive analysis of (prote)omics data’, Nat Methods, vol. 13, no. 9, pp. 731–740, Sep. 2016, doi: 10.1038/nmeth.3901.

[54] J. Cox and M. Mann, ‘1D and 2D annotation enrichment: a statistical method integrating quantitative proteomics with complementary high-throughput data’, BMC Bioinformatics, vol. 13, no. 16, p. S12, Nov. 2012, doi: 10.1186/1471-2105-13-S16-S12.

[55] P. D. Thomas, D. Ebert, A. Muruganujan, T. Mushayahama, L.-P. Albou, and H. Mi, ‘PANTHER: Making genome-scale phylogenetics accessible to all’, Protein Science, vol. 31, no. 1, pp. 8–22, 2022, doi: 10.1002/pro.4218.

[56] M. Ashburner et al., ‘Gene Ontology: tool for the unification of biology’, Nat Genet, vol. 25, no. 1, pp. 25–29, May 2000, doi: 10.1038/75556.

[57] The Gene Ontology Consortium et al., ‘The Gene Ontology knowledgebase in 2023’, Genetics, vol. 224, no. 1, p. iyad031, May 2023, doi: 10.1093/genetics/iyad031.

[58] V. van der Velpen et al., ‘Systemic and central nervous system metabolic alterations in Alzheimer’s disease’, Alzheimers Res Ther, vol. 11, no. 1, p. 93, Nov. 2019, doi: 10.1186/s13195-019-0551-7.

[59] H. Gallart-Ayala et al., ‘A global HILIC-MS approach to measure polar human cerebrospinal fluid metabolome: Exploring gender-associated variation in a cohort of elderly cognitively healthy subjects’, Anal Chim Acta, vol. 1037, pp. 327–337, Dec. 2018, doi: 10.1016/j.aca.2018.04.002.

